# Destabilisation of *bam* transcripts terminates the mitotic phase of *Drosophila* female germline differentiation

**DOI:** 10.1101/2024.07.24.604138

**Authors:** Tamsin J Samuels, Elizabeth J Torley, Emily L Naden, Phoebe E Blair, Frankjel A Hernandez Frometa, Felipe Karam Teixeira

## Abstract

The tight control of the mitotic phase of differentiation is crucial to prevent tumourigenesis while securing tissue homeostasis. In the *Drosophila* female germline, differentiation involves precisely four mitotic divisions, and accumulating evidence suggests that *bag-of-marbles* (*bam*), the initiator of differentiation, is also involved in controlling the number of divisions. To test this hypothesis, we depleted Bam from differentiating cells and found a reduced number of mitotic divisions. We examined the regulation of Bam using RNA imaging methods and found that the *bam* 39 UTR conveys instability to the transcript in the 8-cell cyst and early 16-cell cyst. We show that the RNA binding protein, Rbp9, is responsible for timing *bam* mRNA decay. Rbp9 itself is part of a sequential cascade of RNA binding proteins activated downstream of Bam, and we show that it is regulated through a change in transcription start site, driven by Rbfox1. Altogether, we propose a model in which Bam expression at the dawn of differentiation initiates a series of events that eventually terminates the Bam expression domain.

## Introduction

Adult stem cells divide repeatedly, producing new cells that differentiate to perform specialised functions, maintaining tissue homeostasis. The differentiation process of many adult stem cells includes a phase of transit-amplification, in which mitotic divisions increase the pool of differentiating cells, while minimising the number of divisions of the stem cells themselves (Hsu *et al*, 2014; Watt, 2001; Fuchs *et al*, 2004). Limiting stem cell divisions is thought to minimise the intrinsic risks associated with replication errors and uncontrolled proliferation. For the same reasons, the proliferative transit-amplifying phase of differentiation must be tightly controlled to protect against tumorigenesis.

*Drosophila* female germline stem cells (GSCs), which divide throughout adulthood to produce oocytes, are located in a structure called the germarium at the anterior of the ovary, where they are maintained in a stem cell niche (Fuller & Spradling, 2007). Upon exit from the niche, daughter cells (cystoblasts, CBs) enter differentiation and undergo precisely four mitotic divisions with incomplete cytokinesis to produce a 16-cell cyst (16cc) connected by a structure called the fusome, before terminally differentiating into 16-cell egg chambers. The transition from GSC to CB is initiated by the transcriptional upregulation of Bag-of-marbles (Bam), which is repressed in GSCs by BMP/Dpp signalling from the stem cell niche (Mckearin & Spradling, 1990; McKearin & Ohlstein, 1995; Chen & McKearin, 2003b, 2003a; Song *et al*, 2004). When Bam expression was first observed in the mitotic cells of the germarium, Bam was immediately postulated as part of a mechanism to regulate mitotic division (McKearin & Ohlstein, 1995). However, the essential role of Bam in early differentiation means that *bam* mutant CBs do not enter differentiation and so an additional role for Bam in the exit from mitosis can not be easily examined.

In spite of this challenge, several studies have pointed to a role for Bam in promoting mitotic divisions. *encore* mutants, named due to the phenotype of an additional fifth mitotic division, exhibit an expanded domain of *bam* mRNA (Hawkins *et al*, 1996). Furthermore, while a *bam*^-/-^ mutant can be partially rescued with an inducible heat shock-driven *bam* construct, around 50% of rescued egg chambers were found to contain only 8 cells (i.e. from three mitotic divisions) (Ohlstein & McKearin, 1997). More recently, inducing overexpression of Bam in a *wild type* background was shown to generate 32-cell egg chambers, with a stabilised mutated Bam construct exhibiting a larger effect (Ji *et al*, 2017). Interestingly, overexpression of Cyclin A also leads to 32-cell egg chambers, and this phenotype is partially rescued by reducing the dosage of Bam, which usually stabilises Cyclin A (Lilly *et al*, 2000; Ji *et al*, 2017).

Here, we delve further into the regulation of Bam at the exit of mitosis. We show that depleting Bam in the mitotic region leads to the formation of 8-cell egg chambers. Using single molecule fluorescent *in situ* hybridisation (smFISH) and confocal imaging, we find that *bam* mRNA is rapidly cleared at the 8cc and early 16cc, and that the *bam* 39 UTR is sufficient for this selective destabilisation. Using a series of genetic tools, we show that Rbp9 is required for destabilising *bam* mRNA and restricting the domain of Bam protein expression. Given the central role of Rbp9, we examine its regulation during differentiation to reveal an intricate process involving both translation repression and the non-overlapping use of alternative transcription start sites, with the latter being driven downstream of the cytoplasmic isoform of Rbfox1. We suggest a model in which Bam expression upon exit from the niche initiates a self-limiting clock via Rbfox1 and Rbp9, that eventually leads to Bam downregulation and exit from the mitotic phase of differentiation.

## Results

### Depletion of Bam leads to fewer mitotic divisions during GSC differentiation

It has been shown that ectopic Bam expression leads to an additional mitotic division during GSC differentiation (Ji *et al*, 2017), but it is not clear whether Bam is required for completing the normal four mitotic divisions. Depleting Bam with *bam* RNAi throughout germline development (e.g. with a *nos-GAL4* driver) results in a complete block of differentiation and a tumourous accumulation of GSC-like cells (Blake *et al*, 2017). To deplete Bam from differentiating cells without impacting the initiation of differentiation, we used the *bam-GAL4* driver, which is only activated upon niche exclusion. With *bam-Gal4*, *bam* RNAi is only induced where the *bam* promoter itself is active, in differentiating cells. In agreement with this, we dissected ovaries and observed no tumourous germaria, suggesting that differentiation is initiated normally. However, we found that 47% of ovarioles included at least one 8-cell egg chamber, which was very rarely observed in the *mCherry* RNAi control (**Figure 1A, B**). This result shows that lowering Bam expression reduces the number of mitotic divisions during differentiation, and in combination with the previously published results, we conclude that regulation of the Bam expression domain controls the number of mitotic divisions during female GSC differentiation.

**Figure 1.**
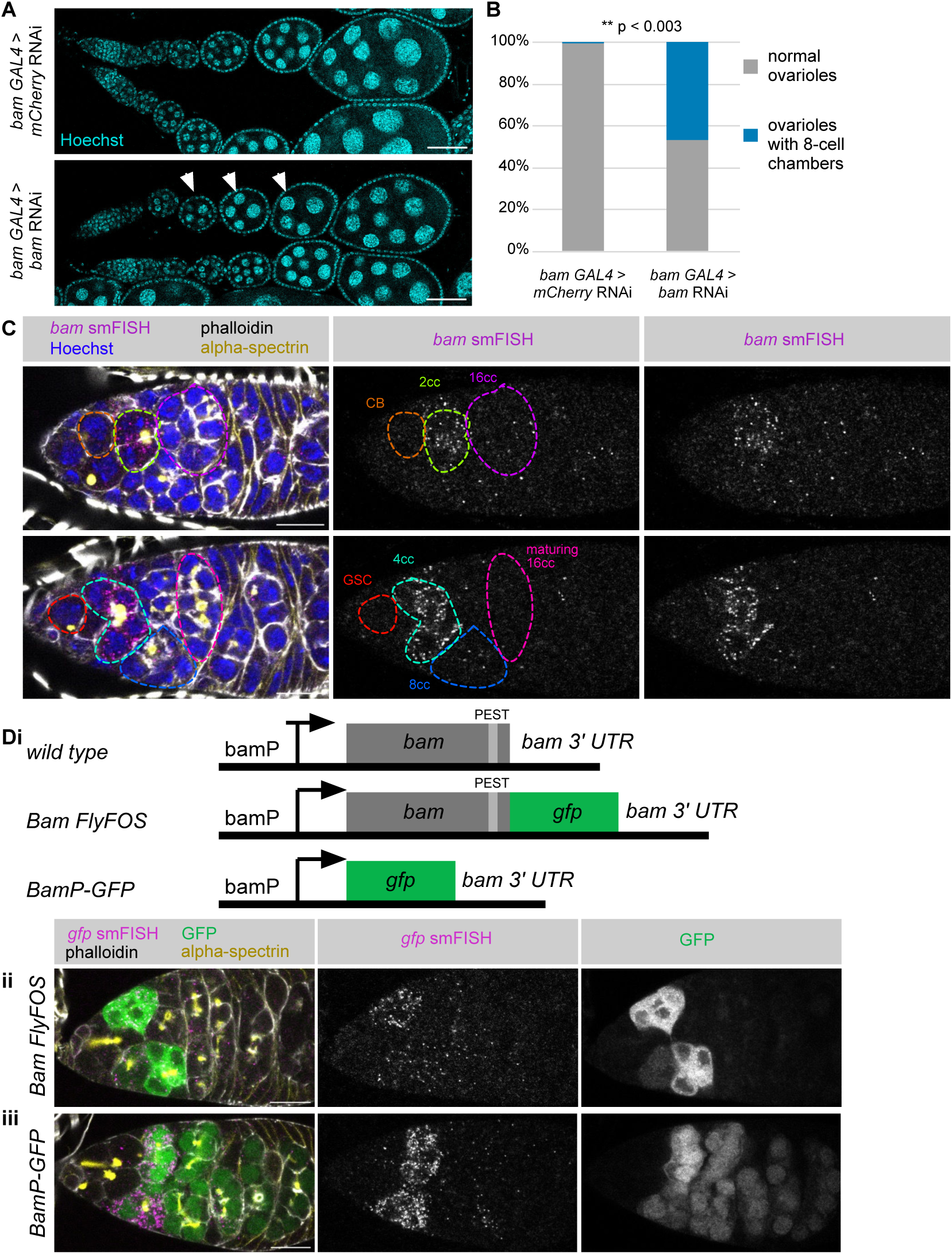
Bam depletion reduces the number of mitotic divisions during differentiation. (**A**) Example ovarioles from *mCherry* RNAi or *bam* RNAi driven by *bam-GAL4*, stained with Hoechst (DNA, blue). White arrows indicate 8 cell egg chambers. Scale bar 50 ¿m. (**B**) Quantitation of the proportion of ovarioles with one or more 8 cell egg chambers (which appear as 4 nuclei in the single z section shown). n = 300 ovarioles for each genotype, from three replicate experiments. p < 0.003, t-test. (**C**) *wild type* germarium stained for *bam* mRNA (smFISH, magenta and gray-scale), f-actin (phalloidin, gray-scale), fusome (alpha-spectrin, yellow) and DNA (Hoechst, blue). Images are two z slices from a single stack. Example GSC and different cyst stages are outlined and labelled. An unannotated gray-scale of the smFISH is also shown. (**Di**) Cartoon depicting the different transgenes used. Staining for GFP protein (green), *gfp* smFISH (magenta), f-actin (phalloidin, gray-scale) and fusome (alpha-spectrin, yellow) in example germaria from (**ii**) *BamFlyFOS* (Sarov *et al*, 2016) (*sfGFP* probes), (**iii**) *BamPGFP (Chen & McKearin, 2003b)* (*eGFP* probes). Scale bars 15 ¿m.

### *bam* mRNA is highest in 4cc and correlates with Bam protein

To explore how Bam protein expression is regulated, we examined *bam* mRNA expression using smFISH. This method is an advance on imaging methodologies previously used to visualise *bam* transcripts, because it can be combined with dyes and antibodies to clearly mark each stage of differentiation. We performed smFISH against *bam,* alongside staining the spectrosome and fusome (alpha-spectrin antibody), cell boundaries (phalloidin - labels F actin) and DNA (Hoechst) (**Figure 1C**, showing two z sections of the same germarium, **S1A**). As expected, we observed very few *bam* transcripts in the GSCs, before *bam* was upregulated to the highest expression in the 4cc. This finding differs slightly from the earlier finding that *bam* is expressed most highly in CBs and 2cc (Mckearin & Spradling, 1990), which is likely explained by our ability to simultaneously stain cell markers. At the 16cc stage very few *bam* transcripts were observed. We rarely observed nuclear *bam* mRNA spots in 16cc and early egg chambers, suggesting that transcription is switched off in late differentiation. Unfortunately, the *bam* introns are too small to design probes to examine transcription directly.

The region of peak *bam* mRNA expression matches the reported pattern of Bam protein expression (McKearin & Ohlstein, 1995). To examine Bam protein and mRNA together, we made use of a *Bam::GFP* transgene line from the FlyFOS collection (Sarov *et al*, 2016), which includes a GFP tag and all of the endogenous regulatory sequence (**Figure 1Di**). We performed smFISH against *gfp* alongside visualising the GFP protein (**Figure 1Dii**) and found that Bam::GFP protein is highest in cells with the highest *gfp* smFISH signal, and is rapidly depleted at later stages of differentiation. While we were unable to devise a reliable protocol to combine *bam* smFISH with anti-Bam antibody staining, the *wild type* samples we imaged were in agreement with the *Bam::GFP* line (**Figure S1B**).

It has been previously reported that the Bam protein contains a highly destabilising PEST sequence (Rogers *et al*, 1986; Mckearin & Spradling, 1990), so to examine the regulatory role of the coding sequence of Bam, we compared the *Bam::GFP FlyFOS* line with the widely used *BamP-GFP* transgene (Chen & McKearin, 2003b). *BamP-GFP* is driven by the *bam* promoter and uses the *bam* 39 UTR, but the coding sequence (CDS) encodes GFP only (**Figure 1Di**). For this construct, smFISH showed a similar *gfp* mRNA expression domain as *Bam::GFP FlyFOS*, but the GFP protein persisted much longer, into the terminally differentiated 16ccs (**Figure 1Diii**). We conclude that the Bam::GFP fusion is less stable than GFP alone, likely due to the PEST sequence in Bam, though other differences in the protein fusions could play a role. The instability of Bam protein means that *bam* mRNA level is the primary determinant of Bam protein level. This finding is supported by our observation that Bam protein levels closely follow *bam* mRNA levels after induction by heat shock and subsequent decay (Samuels *et al*, 2024).

### The *bam* 39 UTR conveys mRNA instability in 8cc and early 16cc

At the later stages of differentiation, *bam* mRNA is downregulated either through switching off transcription or increasing decay of *bam* transcripts. To distinguish between these possibilities, we took advantage of the *Bam UTR sensor* line (Pek *et al*, 2009), in which a ubiquitous tubulin promoter drives a transcript encoding GFP followed by the *bam* 39 UTR (**Figure 2A, S2A**). 39 UTRs contain regulatory sequences, with roles including controlling transcript stability. In ovaries of the *Bam UTR sensor* line, *gfp* transcripts were observed in the GSCs (where the tubulin promoter is active, but the *bam* promoter is not) and throughout differentiation, but with a distinctive absence of cytoplasmic *gfp* transcripts at the 8cc (outlined light blue) and early 16cc (outlined dark blue). Nuclear *gfp* smFISH signal was observed in 8ccs and 16cc (light blue arrows), confirming that the *Bam UTR sensor* transgene is transcriptionally active in these cells, though we could not determine whether the levels of transcription change in these stages. In later 16cc and early egg chambers, *gfp* transcripts were again observed in the cytoplasm. This result suggests that the *gfp* mRNA is specifically unstable in the 8cc and 16cc, via its *bam* 39 UTR. The GFP protein persists into the 8cc and 16cc, likely because the GFP protein is more stable than Bam, as discussed above.

**Figure 2.**
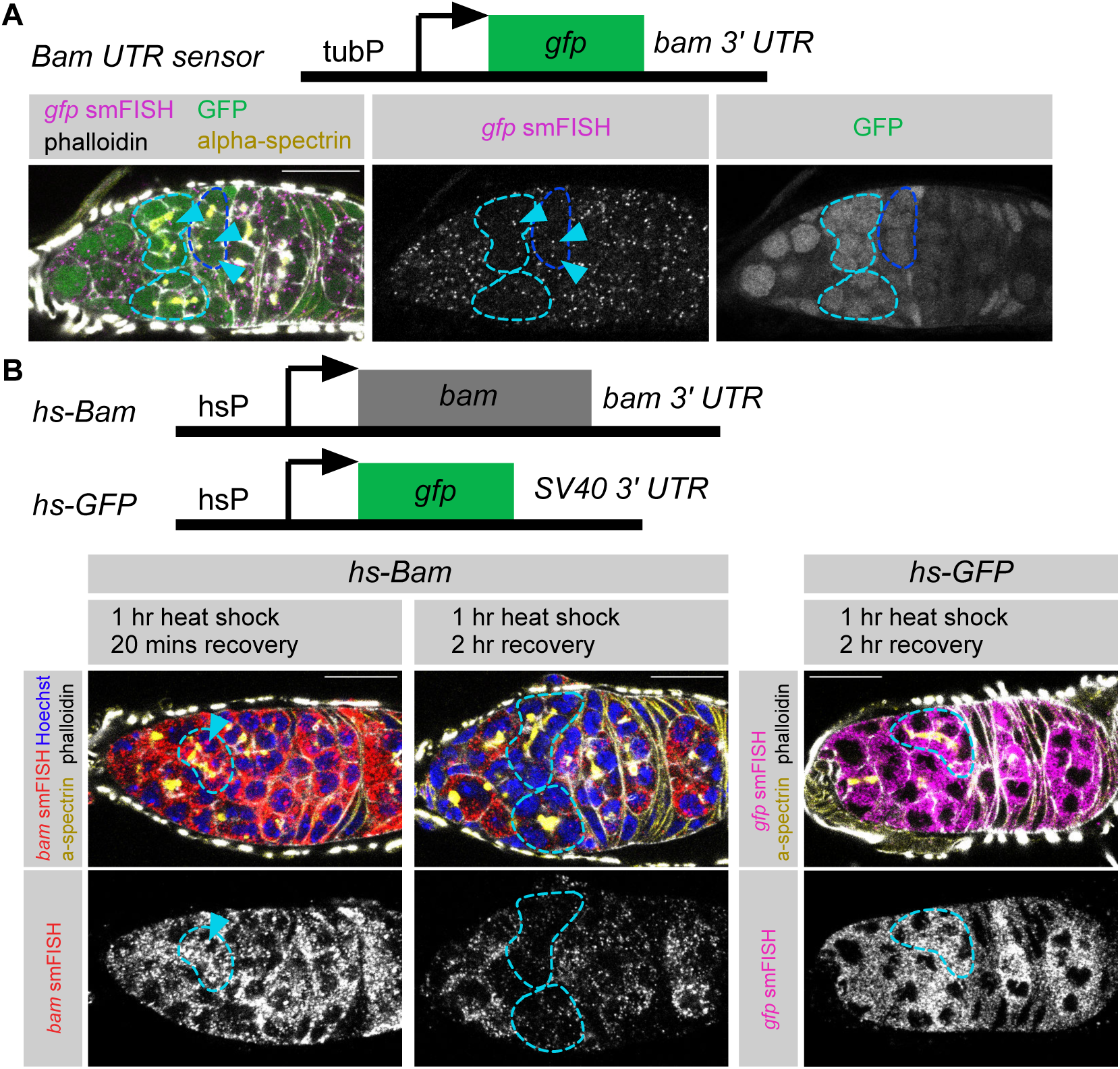
*bam 39 UTR* conveys transcript instability in 8cc and early 16cc. (**A**) Cartoon depicting the *Bam UTR sensor* transgene (Pek *et al*, 2009). Example germaria staining for GFP protein (green), *gfp* smFISH (magenta, *egfp* probes), f-actin (phalloidin, gray-scale) and fusome (alpha-spectrin, yellow). 8cc outlined in light blue, early 16cc outlined in dark blue, nuclear *gfp* smFISH signal indicated with light blue arrow. (**B**) A cartoon depicting the *hs-bam* transgene *(Ohlstein & McKearin, 1997)* and *hs-gfp* transgene. Staining for *bam* smFISH (red and grayscale) or *gfp* smFISH (magenta and grayscale, *egfp* probes), f-actin (phalloidin, gray-scale), fusome (alpha-spectrin, yellow) and DNA (Hoechst, blue) in example germaria after 1 hr 37°C heat shock followed by either 20 mins or 1 hr at 25°C. 8cc outlined in light blue, transcription focus indicated with light blue arrow. Scale bars 15 ¿m.

To examine *bam* mRNA stability using an alternative approach, we performed a 8pulse-chase9 inspired experiment using the *heat-shock-bam* line, in which the Bam CDS followed by the *bam* 39 UTR is transcribed upon heat shock treatment (Ohlstein & McKearin, 1997) (**Figure 2B)**. We performed a 1 hour heat shock (HS) at 37°C, returned the flies to 25°C and then examined *bam* mRNA after 20 minutes and after 2 hours (**Figure 2B, S2B).** Of note, given the strength of the heat-shock promoter, the exogenous *hs-bam*-derived transcripts are expressed at a much higher level than endogenous *bam*. 20 minutes after HS, *bam* was highly expressed in all cells, and bright transcription foci were clearly observed in 8cc (outlined light blue, arrow). However, 2 hours after HS, there was a distinct depletion of *bam* transcripts from the 8ccs (outlined light blue). In later 16cc and early egg chambers, *bam* transcripts were as stable as in the GSC to 4cc domain. This specific transcript depletion at the 8cc stage (outlined light blue) was not observed in a *heat-shock-GFP* control with an *SV40* 39 UTR.

Collectively, these results suggest that *bam* mRNA is unstable at the 8cc and early 16cc stages, and this change in stability is mediated by the *bam* 39 UTR. Interestingly, *bam* is stable in late 16cc and early egg chambers, but in *wild type* ovaries, *bam* transcripts do not accumulate at these stages. This finding suggests that *bam* transcription is eventually switched off upon terminal differentiation, supporting our earlier observation that nuclear *bam* transcription foci are rarely observed at late 16cc and early egg chambers (**Figure 1C**).

### Rbp9 directs *bam* mRNA decay in late differentiation

Since *bam* mRNA is selectively destabilised in the 8cc and early 16cc stages via its 39 UTR, we reasoned that disrupting the RNA decay machinery would stabilise *bam* mRNA and expand the domain of *bam* mRNA expression. Knocking down *pacman* (*pcm*, *xrn1*, an exoribonuclease that degrades decapped RNA) resulted in *bam* mRNA persistence into the 16cc (**Figure 3A,B**), as well as cell death and defects in egg chamber cell number (**Figure S3Ai**). Knocking down *twin* (the CCR4 deadenylase) also expanded the region of *bam* mRNA expression and caused egg chamber cell number defects (**Figure 3C, S3Aii**) (Morris *et al*, 2005; Joly *et al*, 2013). However, while *bam* mRNA is specifically degraded at the 8cc and early 16cc stages, these core mRNA decay components are known to be pervasively expressed (**Figure S3B**) (Samuels *et al*, 2024) and likely degrade many different transcripts in the germline. Therefore, it is most plausible that an intermediate protein acts to recognise *bam* mRNA and direct it to the RNA decay machinery at the correct stage of differentiation.

**Figure 3.**
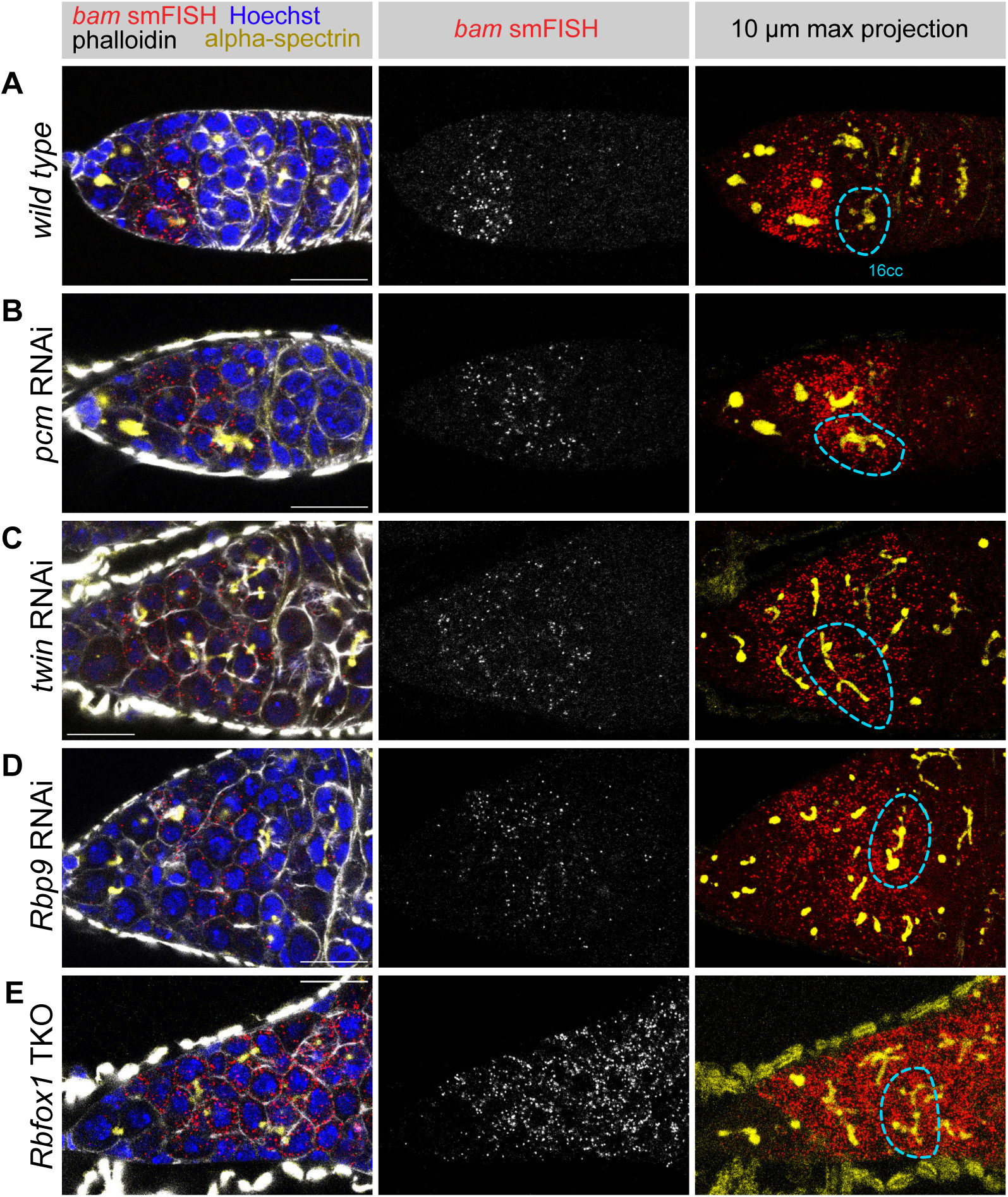
Rbp9 directs *bam* mRNA decay at the end of the proliferative phase. Example germaria from (**A**) *wild type*, and *nos-gal4* (*P{UAS-Dcr-2.D}1; nosP-GAL4-NGT40; nos-GAL4::VP16*) driving (**B**) *pcm* RNAi, (**C**) *twin* RNAi, (**D**) *Rbp9* RNAi and (**E**) germline specific *nos-Cas9* targeting Rbfox1, stained for *bam* mRNA (red and gray-scale), f-actin (phalloidin, gray-scale), fusome (alpha-spectrin, yellow) and DNA (Hoechst, blue). Right hand panel shows a maximum projection of 10 z slices at 1 ¿m spacing, 16cc outlined in light blue Scale bars 15 ¿m.

Rbp9 is a strong candidate for directing *bam* mRNA decay: Rbp9 depletion was previously shown to expand the domain of Bam protein expression, Rbp9 binds the *bam 39 UTR in vitro*, and Rbp9 is upregulated at the 8cc stage (Kim-Ha *et al*, 1999; Jeong & Kim-Ha, 2004). To determine whether Rbp9 downregulates Bam via RNA stability or translation control, we performed smFISH for *bam* mRNA on ovaries of *rbp9* RNAi. In these samples, we observed persistent *bam* mRNA in 8cc and 16cc, beyond the usual stage of *bam* decay (**Figure 3D**). Together, this result suggests that Rbp9 destabilises *bam* mRNA, perhaps by recruiting the RNA decay machinery to the *bam* transcript. It is notable that even in the depletions of *rbp9* and decay machinery, the persistent *bam* mRNA expression domain is eventually downregulated, which is likely explained because *bam* transcription is switched off in later differentiation.

### A cascade of gene regulatory events eventually limits Bam expression

We have shown that Rbp9 is required for the decay of the *bam* mRNA, so we asked how Rbp9 itself is regulated during differentiation. Rbp9 is part of a cascade of RNA binding proteins (RBPs) that are temporally activated during differentiation (Tastan *et al*, 2010). Once Bam is expressed in the CB, cytoplasmic Rbfox1 (previously A2BP1) is upregulated, followed by Rbp9. Both *rbp9* and *rbfox1* mutants exhibit a range of phenotypes from germline cystic tumours to egg chambers with 32 cells (Kim-Ha *et al*, 1999; Tastan *et al*, 2010) (**Figure S3C**). Interestingly, Rbp9 is lost in *rbfox1* mutants and the Bam protein expression domain is expanded (Tastan *et al*, 2010). Similarly, when Rbfox1 was depleted with a germline-specific CRISPR knock out, we found that the region of *bam* mRNA expression was greatly expanded (**Figure 3E**). This is consistent with a model in which Rbfox1 regulates *bam* mRNA via Rbp9.

To examine Rbp9 regulation during differentiation, we performed smFISH and found that *rbp9* mRNA is slightly upregulated during differentiation (**Figure 4A**), with a moderate number of transcripts being reproducibly observed in GSCs and early stages of differentiation when Rbp9 protein is not expressed (Tastan *et al*, 2010). When Rbfox1 was depleted, the accumulation of *rbp9* mRNA was dramatically reduced during differentiation, but the level of *rbp9* transcripts was unchanged in the GSCs (**Figure 4A**).

**Figure 4.**
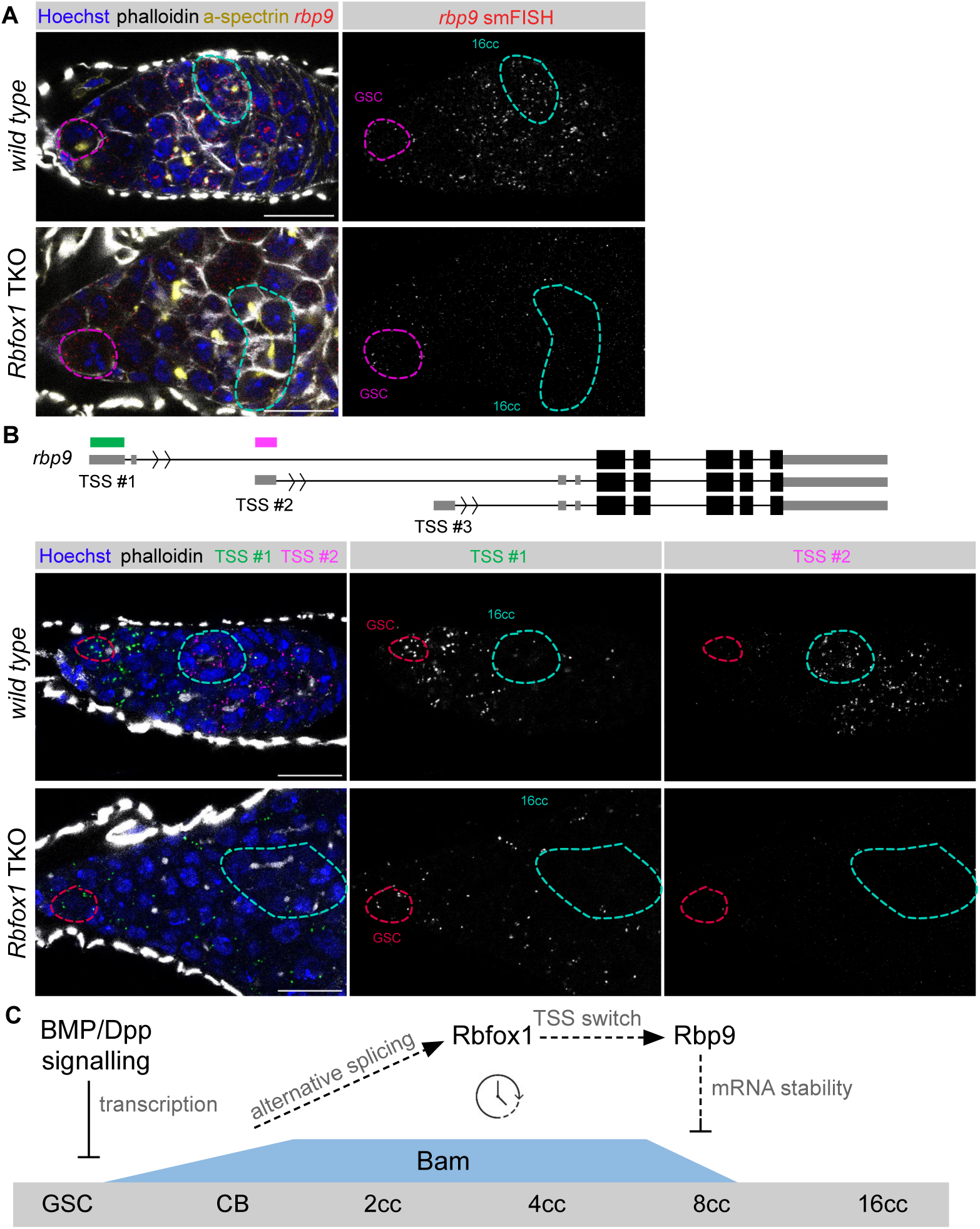
Initiation of a differentiation clock limits Bam expression. (**A**) Example germaria from (**i**) *wild type* and (**ii**) germline specific *nos-Cas9* targeting Rbfox1, stained with smFISH for *rbp9* mRNA (red and gray-scale), f-actin (phalloidin, gray-scale), fusome (alpha-spectrin, yellow) and DNA (Hoechst, blue). GSC and 16cc outlined and labelled. (**B**) Sketch of the *rbp9* gene structure with coloured bars indicating positions of HCR probes. Example germaria from (**i**) *wild type* and (**ii**) germline specific *nos-Cas9* targeting Rbfox1 stained with HCR for *rbp9* TSS #1 (green and gray-scale), *rbp9* TSS #2 (magenta and gray-scale), f-actin (phalloidin, gray-scale), DNA (Hoechst, blue). GSC and 16cc outlined and labelled. (**C**) Proposed model describing a differentiation clock initiated by Bam expression, which limits the proliferative phase of differentiation. Dotted lines indicate speculative or potentially indirect mechanisms. Scale bars 15 ¿m.

The *rbp9* gene is annotated to have three different transcription start sites (TSSs), each producing a transcript with the same CDS, but a different 59 UTR (FlyBase, (Gramates *et al*, 2022)). We previously used a GSC synchronised differentiation system to show that the middle TTS, TTS #2, of *rbp9* is activated only in late differentiation (Samuels *et al*, 2024). TSS #2 has been shown to be repressed by the insulator-binding protein Su(Hw) during egg chamber formation (Soshnev *et al*, 2013). To test *rbp9* TSS usage in *wild type* differentiation, we designed hybridisation chain reaction (HCR) probes against each 59 UTR and observed a near complete switch from TSS #1 to TTS #2 during germline differentiation (**Figure 4B, S4**). Remarkably, the timing of the switch to TSS #2 corresponds to the appearance of Rbp9 protein (Tastan *et al*, 2010), suggesting that only the 59 UTR from TSS #2 allows the translation of Rbp9. TSS #3 did not show cytoplasmic germline transcripts, with some expression in somatic and muscle cells, and some transcription foci in the germline (**Figure S4**). Notably, in this experimental design, signal from a downstream TSS probe may also be picked up in the pre-spliced introns from an upstream TSS.

We asked how loss of Rbfox1 affects *rbp9* transcription by testing the TSS usage in the *Rbfox1* germline-specific knock out. Despite the observation of branched fusomes identifying 8ccs, TSS #2 was never activated during differentiation (**Figure 4B**). It is likely that the loss of the TSS #2 isoform upon Rbfox1 depletion prevents the upregulation of the Rbp9 protein (Tastan *et al*, 2010). Interestingly, in the *rbfox1* depletion, TSS #1 was largely switched off in cysts with branched fusomes (light blue outline), suggesting that the two TSSs are regulated independently and the switching off of TSS #1 does not require Rbfox1.

One step upstream, Rbfox1 itself is upregulated during differentiation via alternative splicing, which leads to the production of an isoform that lacks the nuclear localisation signal (Tastan *et al*, 2010; Carreira-Rosario *et al*, 2016). We previously showed that the introduction of Bam protein in a bam^-/-^ mutant is sufficient to drive this switch (Samuels *et al*, 2024). Altogether, our data, combined with the previously published data, supports our proposed model (**Figure 4C**) in which induction of Bam expression initiates a series of events that eventually terminate the Bam expression domain and the mitotic phase of differentiation. At the onset of differentiation in the CB, BMP-mediated repression of *bam* transcription is released. Bam drives an alternative splicing event to upregulate cytoplasmic Rbfox1, which then upregulates Rbp9 via switching to a downstream TSS, including an alternative 59 UTR allowing Rbp9 translation. Rbp9 then binds to the *bam* mRNA via its 39 UTR to initiate RNA decay, and the intrinsic instability of Bam protein via the PEST sequence limits the domain of Bam protein expression, resulting in cells exiting the transit-amplifying mitotic phase.

Expression of Bam in a synchronised GSC differentiation system drives two sequential waves of gene expression, with the first wave being enriched for genes involved in DNA replication and the cell cycle (Samuels *et al*, 2024). Previously, we suggested that the loss of Bam is crucial for allowing cells to enter into the second wave of gene expression changes. This idea is supported by the findings presented here: Bam must be depleted to complete the mitotic phase of differentiation in the *wild type* female germline. This is in contrast to the findings in males, in which a threshold level of Bam accumulated is required for entry to terminal differentiation (Gönczy *et al*, 1997; Insco *et al*, 2009).

It is unclear from our experiments whether the proposed self-limiting Bam regulatory mechanism is tuned to 8count9 division numbers or to 8time9 the proliferative phase. These possibilities might be distinguished with experiments to manipulate the cell division rate. The proposed self-limiting model for Bam expression involves the regulation of gene expression at multiple levels, including transcription, splicing, mRNA stability, translation control and protein stability. In the 8timing9 model, we speculate that the different regulatory mechanisms could influence the speed of the clock: for example, the pace by which transcriptional upregulation or alternative splicing may result in functional changes is probably slower than that of the regulatory layer involved in RNA or protein decay.

Our model most likely overlooks various unknown intermediate regulators - for example the cytoplasmic RBP Rbfox1 likely mediates the nuclear *rbp9* TSS change indirectly, perhaps via downregulation of Su(Hw) at the 8cc/16cc stage (Soshnev *et al*, 2013). The model is further simplified in its omission of additional downstream targets of each regulator, which may have effects beyond the regulation of Bam. For example, Rbfox1 binds to the translational repressor Bruno (Sugimura & Lilly, 2006; Wang & Lin, 2007; Tastan *et al*, 2010), and destabilises *pumilio* mRNA (Carreira-Rosario *et al*, 2016).

It is unclear how the interactions between regulators may differ between tissues: while Bam is germline-specific, both Rbfox1 and Rbp9 are highly expressed in the *Drosophila* nervous system (modENCODE, (Brown *et al*, 2014)). Rbp9 is closely regulated to the *Drosophila* neuronal RBPs Found in neurons (Fne) and Elav, as well as the human ELAVL/Hu family of proteins, which play diverse roles in RNA regulation including splicing, 39 UTR lengthening and RNA stability (Mulligan & Bicknell, 2023). In both fly and human, this family of RBPs are widely involved in differentiation, emphasising the importance of RNA regulation in stem cells and their progeny (Yao *et al*, 1993; Grassi *et al*, 2019; Kota *et al*, 2021).

It is intriguing that the protein which initiates differentiation must be removed to end the transit-amplifying phase and allow entry into terminal differentiation. We speculate that the self-limiting mechanism regulating the Bam expression domain is protective, preventing the tumourous growth downstream of an uncontrolled proliferative phase caused by unlimited Bam expression. The system is made further robust through a layer of transcriptional regulation of *bam*. Other master differentiation factors are also switched off in mature terminally differentiated cell types, such as Prospero in differentiating *Drosophila* neurons (Spana & Doe, 1995) and MyoD in differentiating muscle (Hinterberger *et al*, 1991), suggesting that this could be a widespread mechanism ensuring unidirectional differentiation and preventing tumourigenesis.

## Acknowledgments

We thank Toshie Kai, the Vienna Drosophila Resource Center, and the Bloomington Drosophila Stock Center for fly reagents, and Eleanor Blake for assistance with the HCR protocol. We also gratefully acknowledge the help of the following facilities: the Department of Genetics Fly Facility, and Dr Ian Clark, Dr Antonina J. Kruppa, Dr Ben Sutcliffe and Dr Jonathan D. Howe from the Department of Genetics Imaging Facility, alongside the Wellcome Trust strategic award (105602/Z/14/Z). TJS is a Herchel Smith Postdoctoral Fellow. ELN was supported by Peterhouse College and the Department of Genetics, University of Cambridge. FKT is a Wellcome Trust and Royal Society Sir Henry Dale Fellow (206257/Z/17/Z) and an EMBO YIP Investigator (5025) and is supported by the Human Frontier Science Program (CDA-00032/2018). For the purpose of Open Access, the author has applied a CC BY public copyright licence to any Author Accepted Manuscript (AAM) version arising from this submission.

## Author contributions

TJS and FKT conceived the idea and designed the experiments. TJS, EJT, ELN, PEB and FAHF performed the experiments. TJS analysed the data. TJS and FKT wrote the manuscript.

## Declaration of Interests

Authors declare no competing interests.

## Materials and Methods

### Resource availability

#### Lead Contact

Further information and requests for resources or reagents should be directed to the lead contact Felipe Karam Teixeira.

#### Data and code availability

This paper does not report original code. Any additional information required to reanalyse the data reported in this paper is available from the lead contact upon request.

### *Drosophila* husbandry and genetics

Unless otherwise stated, stocks and crosses were maintained on standard propionic food at 25°C for experiments. For heat shocks, flies were incubated in pre-warmed vials for 1 hr at 37°C and then transferred to fresh vials at 25°C for either 20 mins or 2 hrs before dissection. The *Drosophila melanogaster* stocks used were:

| Stock | Genotype | Origin |
| --- | --- | --- |
| wild type | *w*[1118] | R. Lehmann lab |
| *nos*-GAL4 | *P*{UAS-*Dcr-2.D*}1; *nos*P-GAL4-NGT40; *nos*-GAL4::VP16 | BDSC #25751 |
| *bam*-GAL4 | *P{UAS-Dcr-2.D}1, w[1118]; Sco/CyO; P{bam-GAL4:VP16}/TM6B* | From E. Bach/Y. Yamashita [YY0274]. |
| *mCherry* RNAi | *y1 sc\* v1 sev21; P{VALIUM20- BDSC RRID: BDSC_35785 mCherry}attP2 (mCherry RNAi)* | BDSC #35785 |
| *bam* RNAi | *y[1] v[1];; P{TRiP.HMS00029}attP2 [bam]* | BDSC #33631 |
| Bam FlyFOS | *y[1] w[\*];; bam-2XTY1-SGFP-V5-preTEV-BLRP-3XFLAG[VK00033]* | VDRC #318001 |
| *BamP*-GFP | *;;P(BamP-GFP)* | Chen & McKearin, 2003 |
| Bam UTR sensor | *tubP-GFP-bam3'UTR/TM3, Sb* | Pek et al., 2009 |
| hs-*bam* | *P{w[+mC]=hs-bam.O}18d, w[1118]* | Ohlstein and McKearin, 1997 |
| hs-GFP | *y[1] ac[1] w[\*]; P{w[+mC]=Hsp70-GFP.204}2MI4/TM3, Sb[1]* | BDSC #51354 |
| *pcm* RNAi | *y[1] sc[\*] v[1] sev[21]; P{y[+t7.7] v[+t1.8]=TRiP.HMS01169}attP2* | BDSC #34690 |
| *twin* RNAi | *y[1] sc[\*] v[1] sev[21]; P{y[+t7.7] v[+t1.8]=TRiP.HMS00493}attP2* | BDSC #32490 |
| *rbp9* RNAi | *y[1] v[1]; P{y[+t7.7] v[+t1.8]=TRiP.GL01167}attP2* | BDSC #42796 |
| *nos* Cas9 | *y[1] M{w[+mC]=nanos-Cas9.P}ZH-2A w[\*]* | BDSC #54591 |
| *rbfox1* TKO | *y[1] v[1]; M{v[+t1.8]=WKO.P5-B10}ZH-86Fb* | BDSC #84213 |

### smFISH and antibody staining

smFISH probes against *bam, egfp, sfgfp* and *rbp9* were designed with the Stellaris Probe Designer (Biosearch Technologies) (Supplementary Table 1). Oligos were labelled with ATTO 633 as described by (Gaspar *et al*, 2017). smFISH and antibody staining was performed as in (Samuels *et al*, 2024): ovaries were dissected in PBS, fixed for 25 mins at room temperature (RT) in 4% formaldehyde in 0.3% PBSTX (0.3% Triton-X in 1X PBS) and then washed 3x 15 mins in 0.3% PBSTX. Samples were transferred to Wash buffer (2X saline sodium citrate (SSC), 10% deionised formamide in nuclease-free water) for 10 mins at RT. Hybridisation buffer (2X SSC, 10% deionised formamide, 20mM vanadyl ribonucleoside complex, 0.1 mg/ml BSA, competitor (1:50 dilution of 5 mg/ml E.coli tRNA and 5 mg/ml salmon sperm ssDNA) in nuclease-free water) was prepared and smFISH probes, primary antibodies and phalloidin (Alexa Fluor 405 or 488 Phalloidin, ThermoFisher Scientific) were added. Ovaries were incubated in this Hybridisation buffer at 37°C overnight in the dark, then washed 3x 15 mins in Wash buffer. Samples were incubated with secondary antibodies in Wash buffer for 2 hrs at RT, then finally washed in Wash buffer, with the addition of Hoechst in one wash step. Finally, samples were mounted in VectaShield mounting media (Vector Laboratories).

For *bam* smFISH with anti-Bam antibody, the protocol was adapted by performing the smFISH without any primary antibodies first in an overnight step, then transferring samples to 0.3% PBSTX. Ovaries were incubated with primary antibody in Block (0.1 mg/ml BSA in 0.3% PBSTX) overnight at 4°C, washed and then incubated with secondary antibody in Block for 2 hr at RT before final washes and mounting.

| Antibody | Concentration | Origin |
| --- | --- | --- |
| Alpha-spectrin (mouse) | 1:100 | DSHB Cat#3A9(323 or M10-2) |
| Bam (mouse) | 1:20 | DSHB bam |
| GFP Booster ATTO 488 | 1:500 | Chromotek (gba488-100) |

### Hybridisation Chain Reaction (HCR)

Split-initiator probes for version 3.0 HCR (Molecular Instruments) were designed according to the parameters described by (Choi *et al*, 2018) (Supplementary Table 1). Ovaries were dissected in PBS and fixed for 25 mins at RT in 4% formaldehyde in 0.2% PBSTX (0.2% Triton-X in 1X PBS). Ovaries were washed with 3x 15 mins in 0.2% PBSTX, then samples were transferred to Hybridisation buffer (Molecular Instruments) for 30 mins at 37°C. 1.6 ¿l of 1¿M probes stock for each probe were added to 100 ¿l of Hybridisation buffer and this was incubated with ovaries at 37°C overnight. On the second day, ovaries were washed with pre-warmed Probe Wash Buffer for 4x 15 mins at 37°C. Samples were then washed 3x 5 mins in 5X SSCT (5X SSC in 0.1% Tween in water) at RT and then left in 5X SSCT until the evening. Samples were incubated in pre-amplification buffer for 30 mins at RT and the hairpins (B2-488, B2-594, B3-647) were prepared according to manufacturer9s instructions (heating to 95°C and then cooling to RT). Samples were incubated with hairpins in amplification buffer overnight in the dark at RT. Ovaries were finally washed in 3x 15 mins in 5X SSCT, with the addition of Hoechst in one wash step. Finally, samples were mounted in VectaShield mounting media (Vector Laboratories).

### Imaging

Images were acquired on a Leica SP8 confocal microscope with a 20x dry objective or 40x oil objective. All imaging experiments were performed for at least 3 replicates. Image processing was performed using Fiji (Schindelin *et al*, 2012).

## Supplementary Figures

**Supplemental Figure 1.**
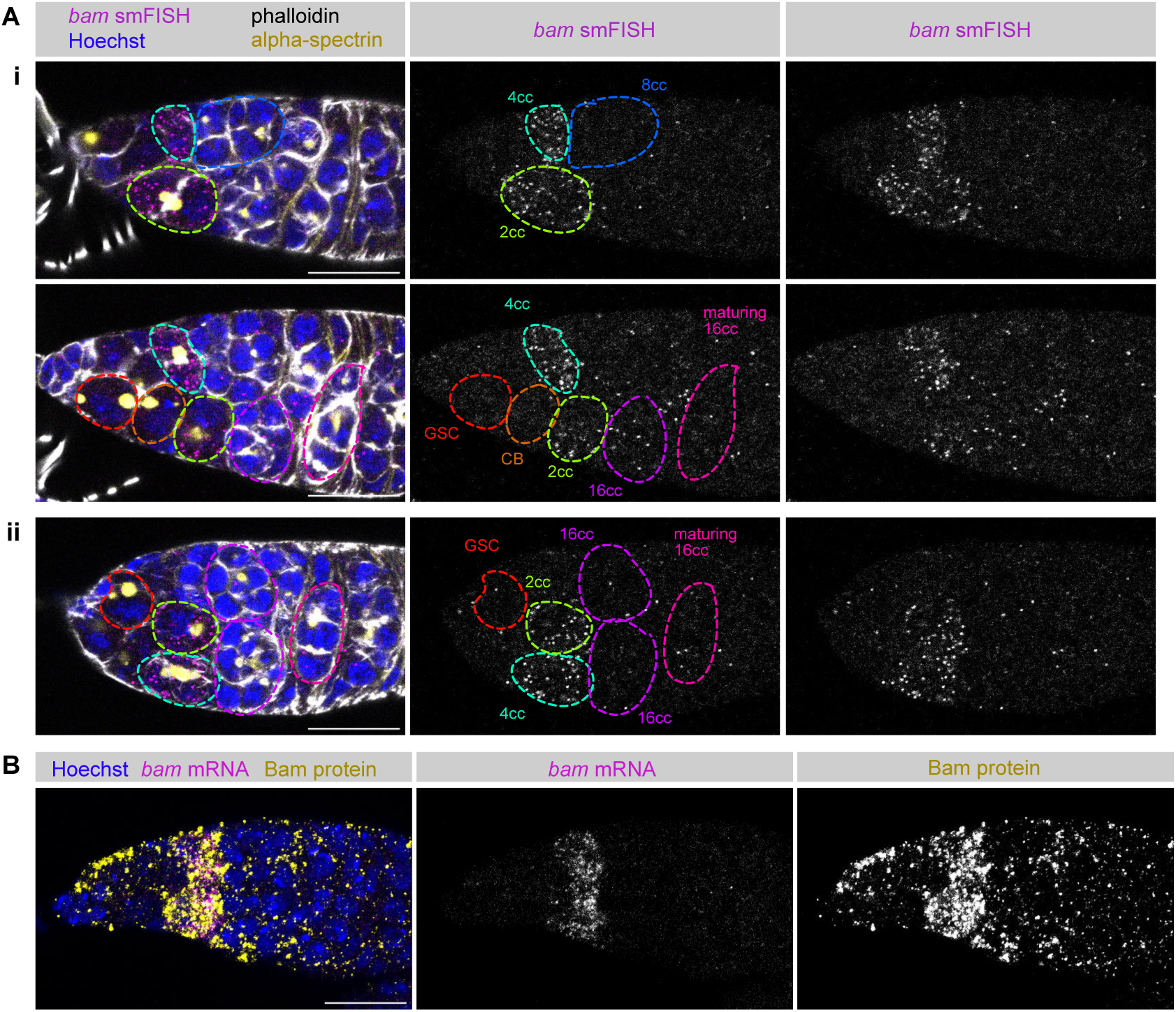
(A) Replicate data of Figure 1C - *wild type* germaria (i, two z sections, and ii are replicates) stained for *bam* mRNA (smFISH, magenta and gray-scale), f-actin (phalloidin, gray-scale), fusome (alpha-spectrin, yellow) and DNA (Hoechst, blue). Example GSC and different stages of cyst are outlined and labelled. An unannotated grayscale of the smFISH is also shown. (**B**) *wild type* germarium stained for *bam* mRNA (smFISH, magenta and gray-scale), Bam protein (yellow and gray-scale) and DNA (Hoechst, blue). Scale bars 15 ¿m.

**Supplemental Figure 2.**
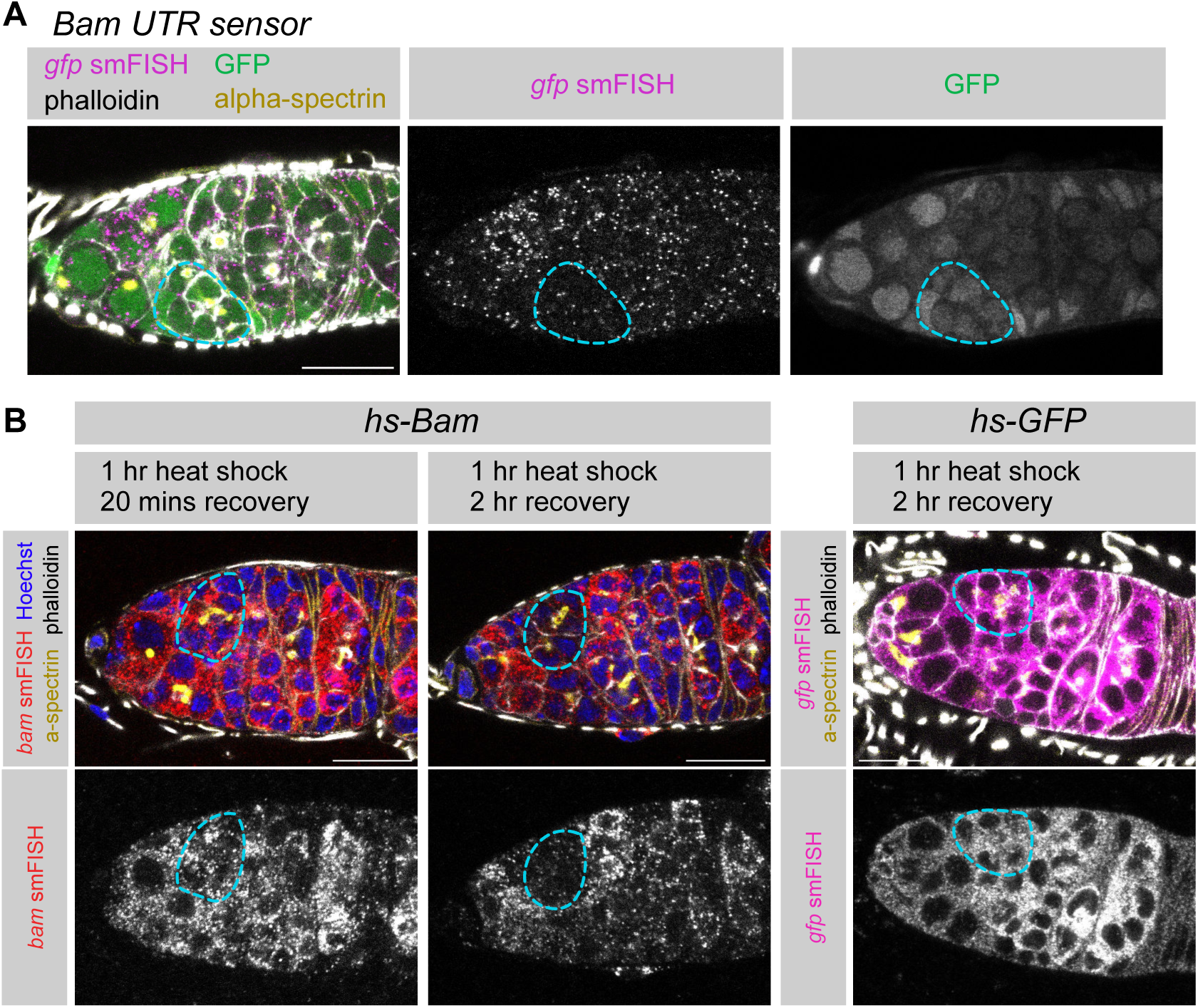
**(A)** Replicate data of Figure 2A 3 example *Bam UTR sensor* germaria staining for GFP protein (green), *gfp* smFISH (magenta, *egfp* probes), f-actin (phalloidin, gray-scale) and fusome (alpha-spectrin, yellow). 8cc outlined in light blue. (**B**) Replicate data of Figure 2B 3 *hs-bam* transgene and *hs-gfp* transgene. Staining for *bam* smFISH (red and grayscale) or *gfp* smFISH (magenta and grayscale, egfp probes), f-actin (phalloidin, gray-scale), fusome (alpha-spectrin, yellow) and DNA (Hoechst, blue) in example germaria after 1 hr 37°C heat shock followed by either 20 mins or 1 hr at 25°C. 8cc outlined in light blue. Scale bars 15

**Supplemental Figure 3.**
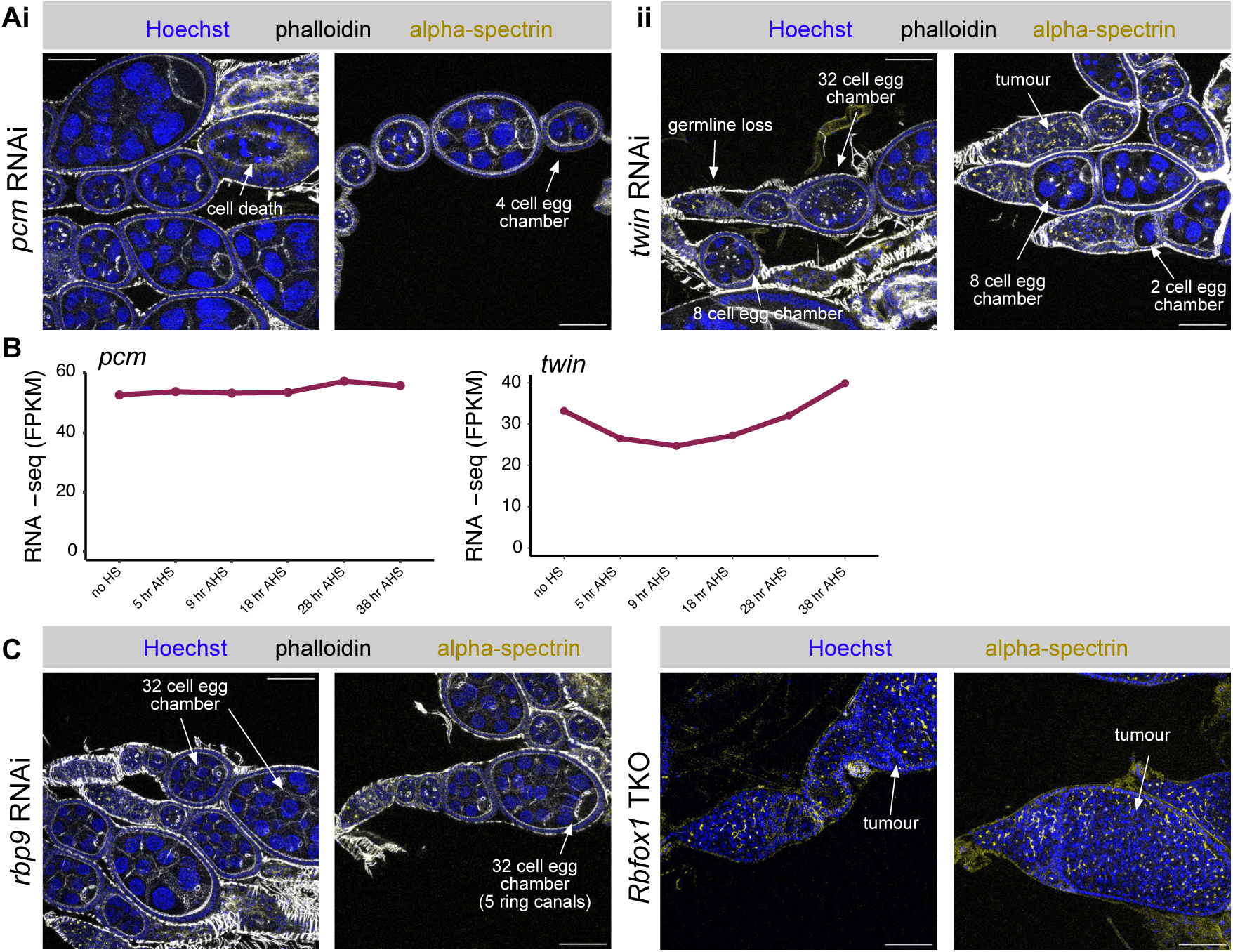
(**A**) Example ovarioles from *nos-gal4* driving (**i**) *pcm* RNAi, (**ii**) *twin* RNAi stained for f-actin (phalloidin, gray-scale), fusome (alpha-spectrin, yellow) and DNA (Hoechst, blue). Images are a maximum projection of 4 z sections at 2 ¿m spacing. (**B**) RNA-seq FPKM (fragments per kilobase of transcript per million reads) for *pcm* and *twin* from synchronised differentiating GSCs over time (Samuels *et al*, 2024). In this system: no heat shock - GSC, 5 hr after heat shock (AHS) - cystoblast, 9 hr AHS - 2cc, 18 hr AHS - 4cc, 28 hr AHS - 8cc, 38 hr AHS - 16cc. (**C**) Example ovarioles from *nos-gal4* driving *Rbp9* RNAi stained for f-actin (phalloidin, gray-scale), fusome (alpha-spectrin, yellow) and DNA (Hoechst, blue) and germline specific *nos-Cas9* targeting Rbfox1, stained for fusome (alpha-spectrin, yellow) and DNA (Hoechst, blue). Images are a maximum projection of 4 z sections at 2 ¿m spacing. Scale bars 50 ¿m.

**Supplemental Figure 4.**
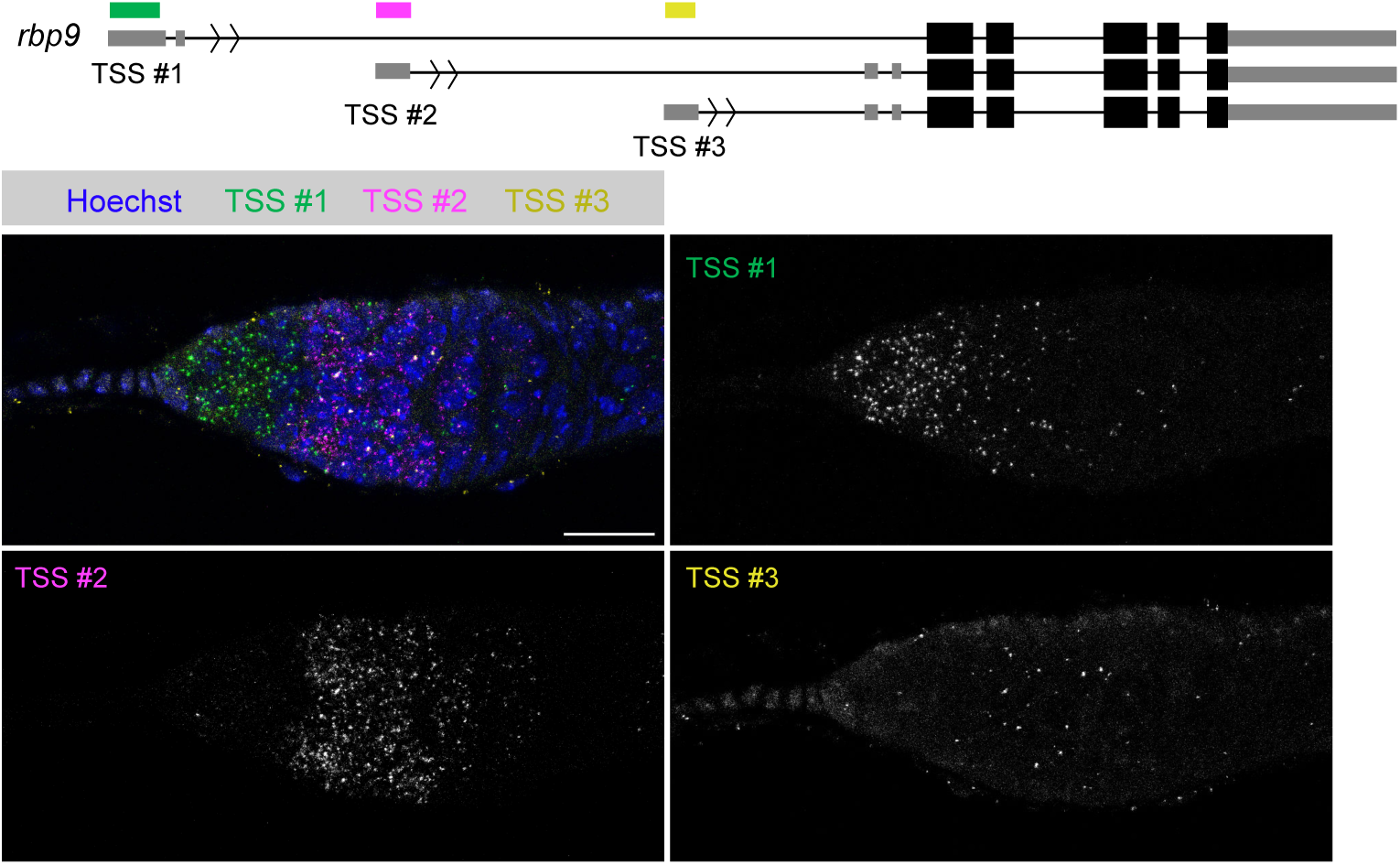
Sketch of the *rbp9* gene structure with coloured bars indicated positions of HCR probes. Example *wild type* germarium stained with HCR for *rbp9* TSS #1 (green and gray-scale), *rbp9* TSS #2 (magenta and gray-scale), *rbp9* TSS #3 (yellow and gray-scale) and DNA (Hoechst, blue). Scale bar 15 ¿m.

**Supplementary Table 1.** Probe sequences for smFISH and HCR

## Notes

### Competing Interest Statement

The authors have declared no competing interest.

### Summary of Updates

The manuscript has been updated after review with Review Commons. As detailed in the response to reviewers, we have so far added new data (replicate images, shown in figures in S1 and S2 in the revised manuscript) and extensive image annotations throughout the original and new figures to render it easier for readers to interpret the data. We have also updated the text to add clarity/explanation, emphasise where statements are speculative, and add interesting discussion points raised by the reviewers.

